# Cell-free translation is more variable than transcription

**DOI:** 10.1101/073841

**Authors:** Fabio Chizzolini, Michele Forlin, Noël Yeh Martín, Giuliano Berloffa, Dario Cecchi, Sheref S. Mansy

**Affiliations:** CIBIo, University of Trento, via Sommarive 9, 38123 Povo (TN), Italy

**Keywords:** cell-free synthetic biology, transcription-translation, IVTT, PURE system, artificial cell

## Abstract

Although RNA synthesis can be reliably controlled with different T7 transcriptional promoters during cell-free gene expression with the PURE system, protein synthesis remains largely unaffected. To better control protein levels, a series of ribosome binding sites (RBS) was investigated. While RBS strength did strongly affect protein synthesis, the RBS sequence could explain less than half of the variability of the data. Protein expression was found to depend on other factors besides the strength of the RBS, including GC content. The complexity of protein synthesis in comparison to RNA synthesis was observed by the higher degree of variability associated with protein expression. This variability was also observed in an *E*. *coli* cell extract-based system. However, the coefficient of variation was larger with *E*. *coli* RNA polymerase than with T7 RNA polymerase, consistent with the increased complexity of *E*. *coli* RNA polymerase.

---

Cell-free expression systems have a wide-range of applications, from experiments designed to gain insight into the workings of biology^1,2^ to the development of new technologies.^3–5^ Although there are several cell-free systems to choose from, none have been well characterized with respect to the influence of different biological parts on system performance. This is a problem, because complex genetic circuitry requires varying degrees of expression of each component. Perhaps the lack of cell-free standardization data reflects an initial bias towards producing maximum amounts of protein from a single gene, just as commercial vectors typically contain sequences, such as the T7 gene 10 leader sequence, to enhance the recombinant expression of protein.^6^ Additionally, when cell-free systems are used to screen genetic constructs for later insertion into bacteria, protein output is often controlled by adjusting the DNA template concentration rather than by controlling gene expression through sequence changes, e.g. by changing the transcriptional promoter or ribosome binding site.^7^ In this case, each gene product is typically encoded on a separate piece of DNA. However, the use of separate pieces of DNA is not suitable for encapsulation in vesicles for the assembly of artificial cells nor ideal for insertion in a genome of a living cell.

Currently, there are several methods that can be used to try to build genetic circuits with desired performance. One method is to create a computational model that can accurately predict gene expression. Such a model may incorporate the kinetic parameters of association, dissociation, and enzymatic activity as well as the influence of RNA folding. Although much progress has been made,^8–11^ several of the parameters governing transcription-translation are still too poorly characterized to build a model that can predict the behavior of a genetic circuit accurately with a high success rate, although the inclusion of transcriptional insulators greatly improves predictability.^12^ Other kinds of computational models do not directly consider the biophysical process of gene expression, but instead attempt to predict gene expression through the summation of the activity individually characterized biological parts. Instead of relying on a computational model, a library of DNA molecules each containing a different mixture of biological parts can be assembled and then screened for activity.^13,14^ One down side of this approach is that only a small subset of all the potential sequences can be practically tested in the laboratory, decreasing the likelihood of identifying a construct with the sought after behavior. Additionally, the assembly and testing of the library of sequences incurs significant cost and time. Both computational and wet lab screening methods frequently depend on the use of biological parts that were previously characterized by assessing the expression of a fluorescent protein. The assumption is that the substitution of the fluorescent protein by the gene of interest will be largely innocuous to system performance. For the time being, a more reasonable approach may be to exploit computational methods to identify a small subset of potential sequences with the desired behavior followed by screening in the laboratory.

Previously we explored the influence of genetic organization^15^ and T7 transcriptional promoter strength^16^ on cell-free expression with the protein synthesis using recombinant elements (PURE)^17^ system. The data revealed that RNA but not protein concentrations were easily tuned with transcriptional promoter strength with a transcription-translation system that depended on T7 RNA polymerase. It thus seemed that the use of different ribosome binding sites would be more amenable to controlling protein synthesis, since the RBS-ribosome complex is initially mediated by known base pairing interactions that directly correlate with the strength of interaction.^18,19^ Although a few reports have exploited the RBS to modulate gene expression *in vitro*,^20,21^ a systematic investigation with the PURE system^17^ is lacking. Here, we investigated the influence of different ribosome binding sites on gene expression with the PURE system. Unlike the strength of the T7 RNA polymerase, the RBS greatly impacted protein synthesis but not in a predictable manner. The activity of the RBS was strongly influenced by the sequence and thus the structure of the mRNA. To ensure that data from different sets of experiments were comparable, the variability in transcription and translation was assessed. Although the noise associated with cell-free expression in compartmentalized (i.e. water-in-oil emulsion droplets, lipid vesicles, or polydimethylsiloxane containers), low copy number systems have been well characterized^2,22,23^ the variability from reactions under bulk, non-compartmentalized conditions have not been reported. We found that the variability in protein synthesis was larger for RNA synthesis than for single gene constructs. The variability increased for cascading reactions, and that a computational model that included parameters for variability helped to screen genetic constructs for a subset with desired activity.

## RESULTS AND DISCUSSION

**Ribosome binding sites strongly but unreliably control gene expression**. 18 different ribosome binding sites were designed and tested (Table S1). The sequence of the ribosome binding site (RBS) was varied according to three variables, including the number of base pair interactions between the ribosome and the RBS, the position of the base pair interactions within the RBS, and the nucleotide composition of the non-base pairing positions within the RBS. 13 of these sequences were generated using a D-criterion optimal statistical design of experiments methodology^24^ using a multiple linear regression model (expressed RFP = number of base pairing interactions with the RBS + location of the base pairing interactions + identity of the non-base paired positions), so that few sequences could be used to efficiently explore the sequence space of the RBS. The remaining five sequences were taken from previously tested constructs.^15,16^

The set of different ribosome binding sites was tested within a genetic construct encoding a red fluorescent protein (RFP) and with a purified, *E*. coli-based transcription-translation system containing T7 RNA polymerase (i.e. the PURE system).^17^ Contrary to what was previously observed with a series of T7 transcriptional promoters,^16^ the set of ribosome binding sites gave a well dispersed distribution of expressed protein concentration (Figure S1). However, a statistical model built with data from the expression of the 13 D-criterion optimal statistically designed sequences explained less than 50% of the variability of the data (adjusted R-squared=0.49, p-value=0.03). Most of the explained variability came from the correlation (Pearson correlation = 0.5) between RFP expression and the number of potential base pairing interactions between the RBS and the ribosome (Figure S2a), whereas the location of the base paired positions within the RBS and the identity of the non-base paired positions had no discernable effect on the expression of RFP. This suggested that something other than ribosome binding was also significantly impacting expression. To determine whether the sequence of the encoded gene influenced the activity of the ribosome binding site, the same set of ribosome binding sites was tested within a construct encoding green fluorescent protein (GFP) (Figure S1). The correlation between the GFP and RFP data was 0.61 for the same set of 13 sequences (Figure 1), indicating that while a significant correlation existed, data from the expression of one protein could not be used to reliably predict the expression of the other protein. The correlation between the expression of GFP to the number of base pairing interactions with the RBS was 0.87 (with respect to to the 13 D-optimal sequences) (Figure S2b), significantly higher than for the expression of RFP. Since the sequences of these two genes greatly differed (48% identity), it seemed likely that the folding of the mRNA or the codon context strongly influenced protein expression, as previously noted.^25–27^ No correlation was found between gene expression and the position of the base pairing interactions within the RBS nor the composition of the non-base pairing positions within the RBS. Taken together, RBS sequences significantly impacted protein expression, as expected, but the effects were not consistent between different genetic sequences, likely reflecting the influences of competing intramolecular base pairing interactions.

**Figure 1.**
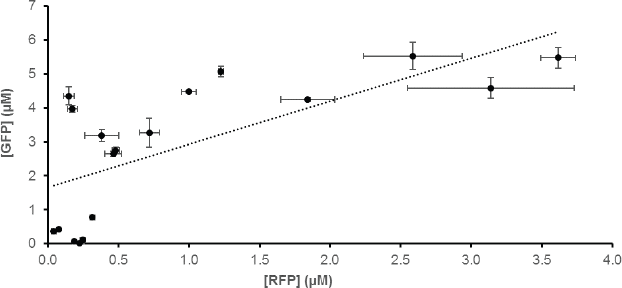
The expression of single gene constructs encoding RFP and GFP do not correlate well. Each point represents a template DNA containing a different ribosome binding site. Reactions were at 37 °C for 6 h. The correlation was 0.61. The different ribosome binding sites employed are listed in Table S1. Sequence details can be found in FC001 (GFP) and FC002 (RFP) of the supporting information.

**The 5’-UTR and 3’-UTR influence protein expression**. To more rapidly screen genetic constructs, linear, PCR product DNA templates were used. Since the incorporation of a transcriptional terminator at the 3’-end of mRNA has been found to increase the expression of protein,^28,29^ we investigated the effect of the 3’-untranslated region (UTR) on the expression of GFP and RFP. The tested DNA templates either ended directly with a stop codon (TAA) or a sequence that encoded a 56 bp sequence of low folding stability, a 91 bp sequence of low folding stability, the bacteriophage T7 Tφ terminator (153 bp), a Spinach aptamer sequence (171 bp), the Tφ terminator embedded in a longer sequence (175 bp), and a sequence containing both the Spinach aptamer and the Tφ terminator (367 bp) (Table S2). Protein expression was the lowest from constructs that lacked a 3’-UTR (Figure S3), as expected.^28,29^ Longer 3’-UTR sequences increased protein yield up to approximately 90 bp in length at which point expression was 26-fold and 12-fold larger for RFP and GFP, respectively, than constructs without a 3’-UTR. There was not a strong dependence on whether the 3’-UTR contained a transcriptional terminator, a random sequence, or an aptamer domain (10% and 17% variance for RFP and GFP, respectively). To confirm that the 3’-UTR was only important for the terminal gene of an operon, two gene operons were constructed encoding both GFP and RFP. The inclusion of the Tφ terminator within the 3’-UTR increased protein synthesis from the second but not the first gene of the operon (Figure S4).

Next, the influence of the 5’-UTR (excluding the ribosome binding site) on gene expression was investigated to identify sequences with similar effects on gene expression to simplify the analysis of other regions of the genetic constructs. Five different 5’-UTR sequences, including four randomly generated sequences lacking strong structural elements (LS1-LS4) and one sequence (LS5) generated by the RBS calculator^30^ were tested (Table S3). Of these five sequences, the different 5’-UTR sequences resulted in different amounts of RNA (based on the fluorescence of the Spinach aptamer) and protein (based on the fluorescence of RFP) synthesis, with three 5’-UTR sequences (LS1, LS2, and LS3) giving more similar protein output (Figure S5). The influence of each of these three 5’-UTR was also similar across different coding sequences, including those of GFP, RFP, and a blue fluorescent protein (BFP). However, the expression of RFP was lower with LS2 than the other fluorescent proteins (e.g. the final RFP concentration was 44% lower than that of GFP with LS2).

**Variability in protein synthesis outweighs variability in RNA synthesis**. Four T7 transcriptional promoters and four ribosome binding sites were then selected to make sixteen different combinations of genetic constructs (Table S4, Table S5). Each DNA template contained a LS1 leader sequence and a Tφ terminator sequence. For both RFP and BFP, the relative transcriptional promoter strength gave predictable distributions of mRNA concentrations regardless of the strength of the RBS (Figure 2a, Figure S6). This is consistent with our previous data on a much larger set of T7 transcriptional promoters with a consistent RBS.^16^ Constructs encoding GFP were not assessed because of spectral overlap between GFP and the encoded Spinach aptamer used to quantify the mRNA. The influence of the RBS on the expressed protein concentration again showed poor correlation between constructs encoding RFP, BFP, and GFP for all four transcriptional promoters (Figure 2b, Figure S7).

**Figure 2.**
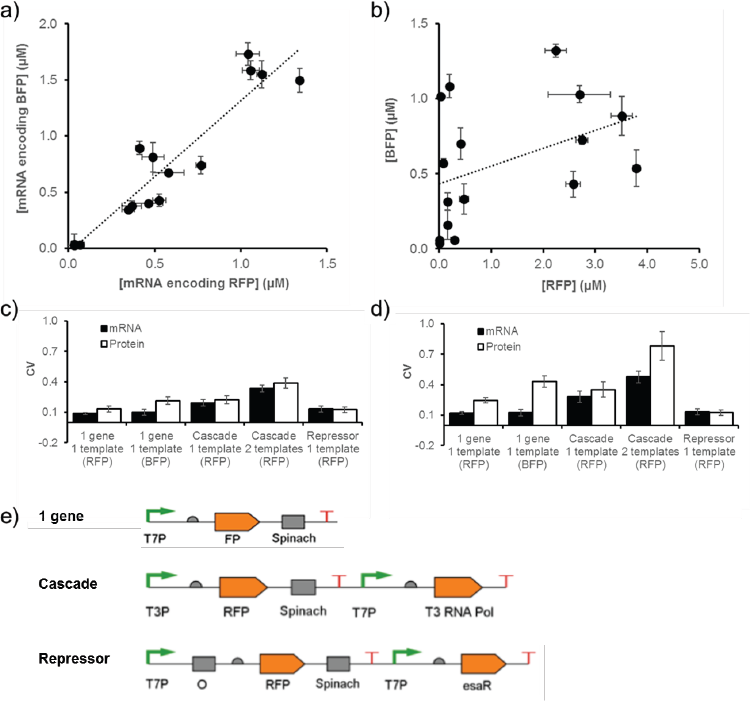
The consistency of transcription and translation in PURE system reactions. (a) Transcriptional promoters predictably control transcription with consistency across different template sequences without influence of the ribosome binding site, whereas (b) the same constructs show significant differences in protein synthesis. Each data point in panels a and b represent the transcription-translation of a different genetic construct with a different combination of transcriptional promoter and RBS. Sequence information and RNA and protein yields can be found in Table S5 and Figure S6. Single (c) and multiple (d) batch transcription-translation reactions show more variability for translation than transcription. The coefficient of variation (CV) was calculated for constructs encoding either one gene or two genes (labeled Cascade and Repressor) for reactions containing only one template or two. All of the templates encoded RFP except for one, which coded for BFP. (e) Single template versions of each genetic construct used in panel c are shown. All of the genetic constructs contained the LS1 5’-UTR for the expression of the reporter gene, i.e. a fluorescent protein followed by Spinach. In the Cascade and Repressor genetic circuits the LS3 5’-UTR was employed for the expression of T3 RNA polymerase and EsaR, respectively. The T7 transcriptional promoter (T7P), T3 transcriptional promoter (T3P), fluorescent protein (FP, representing either RFP or BFP), Spinach aptamer (Spinach), and the binding site for EsaR (O) are all labeled. Sequence details can be found under pFC013A, pDC051A, FC025-FC027, and FC032 in the Inventory of Composable Elements (ICE).

To better assess the variability (or repeatability) of transcription-translation reactions, the coefficient of variation (also called relative standard deviation) was measured for both triplicates run on the same day with transcription-translation machinery from the same batch (total experiments = 3) and on three different days with three different batches of transcriptional-translation machinery (total experiments = 9). The weakest transcriptional promoters were not used for these experiments. The one batch experiments showed larger variability in expressed protein than expressed RNA for both RFP (48% greater variability) and BFP (116% greater variability) encoding constructs (Figure 2c). The multiple batch experiments did not greatly increase the coefficient of variation of transcription, whereas the coefficient of variation of translation increased by 89% for RFP and by 101% for BFP when comparing multiple batch to single batch reactions (Figure 2d). The coefficient of variation of experiments run on the same day from a single batch of transcription-translation machinery reflected, in part, error arising from the assembly of the reaction. The data collected over multiple days were additionally influenced by batch-to-batch variability of transcription-translation solutions. The low coefficient of variation of transcription for both genetic constructs was consistent with the fact that in the PURE system only a single protein is required to synthesize mRNA and that DNA templates possess a consistent structure regardless of sequence, i.e. double stranded DNA typically assumes a b-form α-helix under physiological conditions. Conversely, the increased coefficient of variation observed for translation was consistent with the large number of RNA and protein molecules required to synthesize protein. Further, the folding of the RNA template is sequence dependent and would be expected to affect the efficiency of translation, as observed when comparing RFP and BFP synthesis. In fact, while the coefficient of variation was within 15% for the transcription of RFP and BFP, the coefficient of variation was 65% and 75% greater for the translation of BFP than RFP for one batch and multiple batch experiments, respectively. This is consistent with the lower correlations observed above for the synthesis of different proteins when compared to the synthesis of different mRNA. To determine whether the increase in variability for transcription was dependent on the RNA polymerase or reflected a specific feature arising from the PURE system, we repeated the measurements with T7 and *E. coli* transcriptional promoters in an *E. coli* cell extract. The coefficient of variation was 6-fold larger for transcription than for translation with T7 RNA polymerase, and 1.9-fold larger with *E. coli* RNA polymerase (Figure S8). The coefficient of variation was 10.4-fold larger for transcription with *E. coli* RNA polymerase than with T7 RNA polymerase, consistent with the fact that *E. coli* RNA polymerase is composed of multiple, separately encoded protein subunits.

The variability of transcription-translation was confirmed by assembling transcription-translation reactions in a 1536-well plate to construct a pixelated image. In this way, the clarity of the image would visually reveal the variability of RNA (based on the fluorescence of the Spinach aptamer) and protein (based on the fluorescence of RFP) synthesis. A 196 pixel version of the yin-yang symbol was designed by exploiting genetic constructs that would give rise to four different color intensities based on RNA and protein synthesis levels. Some of the wells that contained the same genetic construct also contained different batches of transcription-translation machinery. Since each color intensity was represented by at least 57 contiguous pixels, variability was easily observed visually. Consistent with the data presented above, the RNA-based image was much clearer than the protein-based image (Figure S9), confirming that RNA synthesis was less variable than protein synthesis.

**A simple computational model that includes parameters for variability can reasonably predict cell-free expression**. A generalization of the computational model described by Stogbauer et al.^8^ was built to serve as a predictive tool for the design of more complex genetic circuits in cell-free systems (Figure S10). In this model, the resources needed for transcription and translation were grouped separately. However, parameters for the strength of the T7 transcriptional promoter and the RBS were also included.^16^ Additionally, two noise parameters were incorporated. The first noise parameter was meant to address the error that was observed from the single batch experiments by allowing the DNA template concentration to fluctuate by ±5%. The second noise parameter approximated the error observed from the multiple batch experiments by allowing the concentrations of components of the PURE system to vary by ±10%. Kinetic parameters were inferred from a subset of experiments with different combinations of T7 transcriptional promoters and RBS sequences (Figure S7). To address the absolute difference in RFP, GFP, and BFP expression, the parameters for the kinetics of transcription and translation (r_rna___prod_ and r_prot___prod_, respectively), as well as the parameter describing protein maturation (r_prot___mat_) were inferred for each protein. The final inferred parameters, along with the starting concentrations of the molecular components and the parameters accounting for noise were also included (Table S6, Table S7, Table S8). The resulting model fit well the experimental data (Figure S11).

Since a wide-range of colors can be made by mixing different intensities of red, blue, and green colors together, we wondered if genetically encoded colored pictures could be produced by expressing different amounts of RFP, BFP, and GFP or if the variability in protein expression would interfere with the predictable formation of new colors. The kinetic model was used to identify genetic constructs that could be assembled into a 25 pixel RGB color triangle. Each well contained three different DNA templates at the same concentration, each encoding a different fluorescent protein. The model was used to indicate which T7 transcriptional promoter and which *E. coli* ribosome binding site could be used to produce 25 different colors. As seen in Figure 3, the assembled reactions produced a color triangle that closely matched the prediction, and the variability associated with the expression of three different proteins was low enough to allow for the development of the image. The average absolute difference between the predictions and the experimental proportions for the three fluorescent proteins was 32%, 11%, and 17% for RFP, GFP, and BFP, respectively. The largest difference between the prediction and the measured protein expression was associated with RFP, which was overestimated by the model. Under the experimental conditions employed, photobleaching did not significantly impact the measurements (Figure S12).

**Figure 3.**
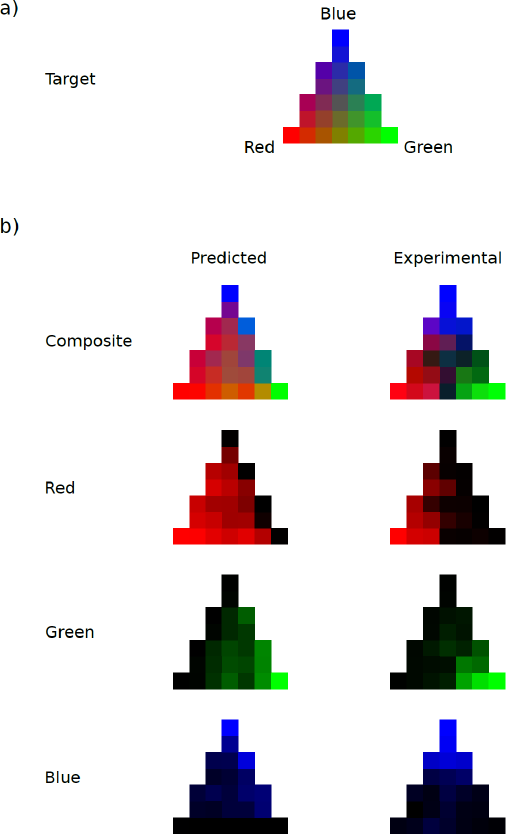
Transcription-translation reactions can be used to construct a RGB color triangle. (a) The target picture is composed of 25 pixels, each displaying a different color resulting from the three different gradients that stem from the vertexes of the triangle. (b) The computational model was employed to decide which combination of promoters and ribosome binding sites should be used to achieve the desired color for each pixel (Figure S14, Table S12). Based on the computational prediction for the expression of each gene in each pixel, a predicted image was generated. Each sequence was then assembled and run.

**Genetic cascades increase the variability of gene expression**. We then explored the effect of a simple genetic cascade on the variability of protein expression. This cascade consisted of the expression of T3 RNA polymerase from a T7 transcriptional promoter followed by the expression of RFP from a T3 transcriptional promoter (Figure 2e). The computational model was updated to include the new parameters (e.g. the kinetics of T3 RNA polymerase) (Table S9, Table S10, Table S11). Thirteen different constructs containing three different T7 transcriptional promoters and three different ribosome binding sites were selected. The model predicted that increased expression of RFP was mainly due to two elements, a medium to weak strength T7 transcriptional promoter and a strong RBS for RFP (Figure S13). By modifying the noise parameters according to their estimated probability distribution, the model did not predict a significant increase in the coefficient of variation for the synthesis of protein in the cascade reaction with respect to the expression of the single gene. The thirteen constructs were then tested in the laboratory in two different ways. The two genes of the cascade were either placed on separate pieces of DNA or together on the same DNA template. Most cell-free genetic circuitry reported in the literature exploit separate pieces of DNA to speed up the screening of biological parts; however, the effect of such architecture on expression and variability are unexplored. Contrary to what the model predicted, the coefficient of variation increased by 120% and 70% for RNA and protein synthesis, respectively, for the expression of the cascade in comparison to that of a single gene for reactions run with a single batch of transcription-translation machinery and using a single DNA template (Figure 2c). The analogous increase in the coefficient of variation for experiments run with multiple batches was 130% and 43% for RNA and protein synthesis, respectively. The fact that the coefficient of variation significantly increased when going from a single gene construct to a simple, two-step cascade suggests that genetic circuitry of increased complexity will concomitantly increase the variability of expression. Also, unlike the single gene data, this specific cascade showed variability in the expression of the final RNA of the cascade that was similar to that of the final protein. This is likely because the first gene of the cascade expressed the RNA polymerase needed to transcribe the second gene. In other words, the variability associated with producing the T3 RNA polymerase directly impacted the extent of transcription of the second gene. The coefficient of variation of the cascade consisting of two pieces of DNA run with multiple batches of transcription-translation machinery was large, so care should be taken in screening genetic circuits with multiple pieces of DNA *in vitro*. The increase in variability of expressed RNA and protein from the cascade was visually confirmed by the construction of a yin-yang image (Figure S15). Finally, the genetic cascade also revealed the existence of the influence of depleting resources. Transcription and translation share common reactants,^9^ and so some degree of competition for resources was expected although not previously observed for non-cascading reactions.^16^ For example, the use of strong transcriptional promoters and ribosome binding sites resulted in relatively low amounts of RFP (Figure S16). The configuration that produced the most RFP contained a weak T7 Transcriptional promoter, a medium strength RBS for T3 RNA polymerase, and a strong RBS for RFP, which was consistent with the kinetic model for the cascade.

Since the inclusion of a transcriptional repressor can be exploited to confer sensing capability on artificial cells^5,31^ and potentially on cascading cell-free networks, we next sought to investigate the behavior of genetic constructs encoding a transcriptional repressor. This construct contained two T7 transcriptional promoters and two ribosome binding sites (Figure 2e). Each transcriptional promoter and RBS pair controlled the expression of a single gene, either RFP or the transcriptional repressor. The region upstream of the RFP encoding sequence also contained an operator sequence to bind the transcriptional repressor. A LuxR homologue, D91G EsaR^32^ (hereafter referred to as EsaR), from the organism *Pantoea stewartii* was used as the transcriptional repressor. In the absence of EsaR’s cognate ligand, EsaR blocks transcription, and in the presence of the quorum signal 3-oxohexanoyl-homoserine lactone (3OC6HSL), expression is derepressed.

After screening a small set of different spacer sequences between the transcriptional promoter and the operator sequence (Table S13, Figure S17), a kinetic model for protein expression under the control of EsaR was built and estimated (Table S14, Table S15, Table S16). Eight different genetic constructs that the model predicted to show a gradient of expression between repressed and derepressed states were assembled onto a single DNA (Table S17). Of these genetic constructs, four showed at least 5-fold derepression of transcription in the presence of 3OC6HSL, and four gave at least 2-fold more cell-free expressed protein when 3OC6HSL was added to the reaction (Figure S18). The transcription and translation data correlated well with the predictions from the computational model (0.74 and 0.87 for RNA and RFP expression, respectively) even if the model predicted higher absolute concentrations. When rescaled to the actual observed concentrations, the expression ratios matched well the experimental data (Figure S19). However, since little RNA is needed to make much protein,^16^ none of the constructs yielded low RFP concentration in the absence of 3OC6HSL and high RFP concentration in the presence of 3OC6HSL. Altering the DNA template concentration from 1.6 nM to 0.42 nM did not significantly impact the difference between repressed and derepressed states (Figure S20). Nevertheless, clear on-off switching activity was observed. The coefficient of variation of expressed protein was similar to the transcription-translation of single gene constructs (Figure 2). The difference was again with the RNA, which was similar to that of the protein. This similarity in variability in expressed RNA and protein was observed with both genetic cascades, the cascade in which the first gene encoded a RNA polymerase and the cascade in which the first gene encoded a transcriptional repressor. In both cases, the variability in the synthesis of the product of the first gene directly impacted the extent of transcription of the second gene.

**A better understanding of the influence of RNA folding on transcription-translation may improve predictability**. It has often been argued that biological parts need to be thoroughly characterized so that a set of rules and equations can be formulated that would allow for the construction of more complex systems with predictable behavior.^33,34^ However, to date this approach has not been successful *in vivo* or *in vitro*. Even in the simplified case of the cell-free transcription-translation of a single gene, the influence of the RBS is not consistent. The inconsistency in the activity of the RBS likely reflects a dependence on the entire sequence of the mRNA. That is, different sequences fold differently which affects the accessibility of the RBS, the initiation of translation,^35^ and ribosome density.^36^ To better probe whether RNA folding impacted the cell-free expression of protein, four different genetic sequences encoding BFP were tested. The original BFP sequence, which was codon optimized for yeast and bacteria, was resynthesized to maintain the same amino acid sequence but to have maximal GC content, minimal GC content, and an additional sequence was made that was specifically codon optimized for *E. coli*. From these four sequences, the GC content appeared to strongly impact the amount of synthesized protein but not the RNA concentration (Figure S21).

To isolate the effect on translation, the protein concentration was divided by the RNA concentration, revealing a clear correlation between higher GC content with the increased expression of protein (Figure 4). That data are similar to what has been observed previously *in vivo* in *Saccharomyces cerevisiae*.^37^ The mechanism of this effect is not known, but increased GC content is associated with an increased density of ribosomes on the mRNA. Although similar studies have not been performed with bacteria, a bias for higher GC content codons is found across all three Kingdoms of life.^38^ More work is needed to determine how robust the relationship between GC content and protein expression is with bacterial systems. Nevertheless, it does appear that mRNA folding significantly affects the expression of protein. Taken together with previous studies on the initiation of translation,^39^ high protein expression may be best achieved with sequences of low structure around the RBS, including the 13 codons after the start codon^35^ and high GC content throughout the remainder of the gene.

**Figure 4.**
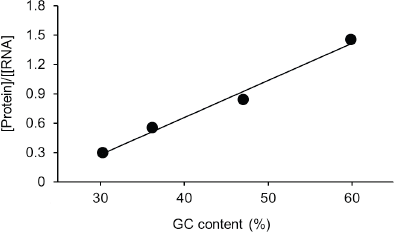
The GC content of the coding sequence of BFP affects the synthesis of protein. The coding sequence of the protein was modified to either maximize the AT content or to maximize the GC content. Additionally, a codon optimized version of the coding sequence was generated in order to assess the impact on translation. Each construct was expressed with the PURE system. The DNA constructs were, from lower GC content to higher GC content, NG046 (AT rich, GC content: 30.4%), NG045 (original sequence, GC content: 36.3%), NG047 (codon optimized, GC content: 47.1%), and NG048 (GC rich, GC content: 60.0%). R-squared = 0.985.

The RBS is not the only complicating factor in predicting the expression of protein. Variability is always present due to the intrinsic stochastic nature of basic molecular events, such as transcription and translation. The stochasticity of compartmentalized systems has been evaluated,^22,23,40^ but the variability of bulk transcription-translation reactions have not been thoroughly investigated. Some of the noise observed in encapsulated, cell-free gene expression was thought to either arise from differences in DNA template copy number across different droplets or from the intrinsic stochasticity of transcription itself.^22,23^ In bulk, where complications arising from encapsulation do not exist, transcription was not found to exhibit much variability. However, variability in transcription significantly increased when different genes of a network were encoded within different strands of DNA. Having genes on separate DNA strands likely introduced variability in the ratio of templates in a somewhat similar way to the influence of DNA template copy number on transcription encapsulated within droplets. In bulk reactions, protein synthesis provided the largest source of variability, which likely reflected, in part, the the complexity of translation and the consequences of using a template that can assume a variety of structures with different degrees of stability. Further, the variability associated with translation is confounded by the batch-to-batch variability of the transcription-translation machinery used to mediate gene expression.

Current computational models do not adequately take into consideration RNA folding. Thus far, models are built by incorporating a data set containing the activity of biological parts, and these resulting models can usually describe accurately the behavior of those same biological parts under previously tested conditions. But much more is needed if these models are to be of real engineering value. It is not sufficient to describe and understand a system. Instead, within a certain margin of error, the behavior of a new device, potentially under new conditions needs to be reliably predicted. Otherwise, the construction of new genetic devices remains a process of trial and error. Unfortunately, the development of a model that fits the experimentally available data does not necessarily mean that the model is capable of predicting experiments that have yet to be run, i.e. there is a difference between model estimation and prediction. For example, the model used herein can predict the expression of each protein individually but the data set for the expression of RFP cannot be used to reliably predict the expression of GFP. It seems that what is missing is a deeper understanding of the role of RNA. A step forward in this direction was the development of an equilibrium statistical thermodynamic model to predict the activity of a RBS sequence within the context of the surrounding RNA.^41^ However, the dynamics governing the folding of mRNA and the relationship between the kinetics of mRNA folding and gene expression are still not sufficiently understood.^42^ Therefore, as noted before,^43,44^ there is approximately a 50% chance that the predictions will fall within two-fold of experimentally determined protein levels. For now, it may be wise to simply exploit tools that incorporate the folding of the RNA, as in the RBS calculator, and the variability in gene expression to identify a restricted set of candidate sequences for screening in the laboratory.

## METHODS

**Genetic Constructs**. Genes encoding the fluorescent proteins GFPmut3b and mRFP1 and the spinach aptamer were from previously described constructs.^16^ Genetic sequence of the T3 RNA polymerase (BBa_K346000) was from the registry of standard biological parts (http://partsregistry.org). Genes encoding the proteins Azurite (referred to as BFP) and D91G EsaR were from Addgene (plasmid #14034 and plasmid #47646, respectively). All other genes were synthesized by Eurofins Scientific. All genes were subcloned into pET21 b by isothermal Gibson assembly.^45^ All constructs were confirmed by sequencing by GATC Biotech.

**Cell-Free Transcription-Translation**: Unless otherwise indicated, 9 μL transcription-translation reactions with the PURExpress in vitro protein synthesis kit (New England Biolabs) contained 12.6 nM of linear DNA template and 4 units of human placenta RNase inhibitor (New England BioLabs). When needed, DFHBI (Lucerna) was added to a final concentration of 60 μM. The reaction components were assembled in an ice-cold metal plate, and the reaction initiated by incubation at 37 °C. Reactions were monitored for 6 h with a Rotor-Gene Q 6plex system (Qiagen). The blue channel was used to detect Azurite (excitation, 365 ±20 nm; emission, 460 ±20 nm), the green channel was used to detect GFPmut3b and Spinach (excitation, 470 ±10 nm; emission, 510 ±5 nm), and the orange channel was used to detect mRFP1 (excitation, 585 ±5 nm; emission, 610 ±5 nm). Each reaction was repeated at least three times. The template DNA concentration was 0.5 nM for single gene expression experiments. The cascade genetic circuit with the two genes on two separate pieces of DNA employed the following template concentrations: 12.6 nM (reporter gene encoding mRFP1 and Spinach) and 4.2 nM (gene coding for T3 RNA polymerase). The cascade genetic circuit with two genes on the same piece of DNA employed a template DNA concentration of 5 nM. Finally, the experiments performed with the repressible genetic circuit employed a template DNA concentration of 5 nM. When used, the concentration of 3-oxohexanoyl-homoserine lactone was 5 μM.

**Cell-free extract preparation**: *E. coli* BL21-Rosetta2(DE3) were grown in in 2xYTP media supplemented with 40 mM K_2_HPO_4_, 22 mM KH_2_PO_4_, and 34 μg/mL chloramphenicol overnight at 37 °C, 220 rpm. Next, an aliquot was diluted to OD^600 nm^ = 0.01 and incubated for 3.5 h at 37 °C, 220 rpm. Cells were then pelleted at 6000 g for 6 min at 4 °C. The extract was then prepared following closely the protocol from the Noireaux laboratory,^46^with the following exceptions. Snakeskin dialysis membranes (10 kDa MWCO Thermo fischer-68100) were used in place of the Slide-A-Lyzer Dialysis Cassettes for 3 h at 4 °C. Aliquots of freshly prepared extract were flash-frozen with liquid nitrogen and stored at -80 °C. The amino acid solution was prepared following the indications of Cashera et al. for equimolar concentration at pH 7.86.^47^ The composition of the cell-free reaction was 10 nM plasmid DNA, 33% (v/v) cell-free extract, 7.14% (v/v) energetic solution, 1.5 mM amino acid solution, 2% (v/v) PEG-8000. The reactions were supplemented with 12 mM maltose for ATP regeneration^48^ and 5 mM or 10 mM Mg^2+^-glutamate for reactions containing T7 RNA polymerase^49^ and the endogenous *E. coli* RNA polymerase, respectively. To monitor mRNA through the encoded Spinach aptamer, reactions contained 60 μM DFHBI (Lucerna), resulting in a final DMSO concentration of 0.4% (v/v).

**Genetically encoded pictures**: 14×14 pixel pictures were generated with four different DNA templates. Following the target picture, the number of different 9 μL PURE system reactions required to fill the pixels was calculated. Reactions were setup with components from different batches of the PURE system. After 6 h of incubation at 37 °C, each reaction was pipetted into the wells (of a ViewPlate-1536 F from Perkin Elmer) of the plate to produce the pixelated image. After filling all of the required wells, the plate was centrifuged at 4000 rpm for 1 min at 4 °C with a Thermo Scientific Legend X1R centrifuge with a T20 microplate rotor. A Typhoon Trio from GE Healthcare was used to visualize the picture. For the RNA picture, a blue laser was used (488 nm) for excitation in combination with a 526 SP filter (short-pass filter transmitting light below 526 nm) for emission. For the protein picture, a green laser was used (532 nm) for excitation in combination with the 610 BP 30 filter (transmitting light between 595 nm and 625 nm) for emission. For both of the pictures, the gain was set to 1000 V.

For the multicolored triangle picture, a 384-well plate (Greiner Bio-one 384 Flat Bottom Black) was employed. Each well, representing a single pixel of the picture, was filled with a 9 μL PURE system reaction. Each reaction included three different DNA templates encoding the fluorescent proteins mRFP1, GFPmut3b, and Azurite, each controlled by the appropriate combination of promoter and ribosome binding site. The reaction was incubated in a PCR tube at 37 °C for 6 h, after which each reaction was placed into the corresponding well of the 384-well plate. Next, a plate reader Tecan Infinite 200 was used to record the fluorescence of Azurite (λ_ex_= 377 nm, λ_em_= 472nm), GFPmut3b (λ_ex_= 474 nm, λ_em_= 511 nm), and mRFP (λ_ex_= 579 nm, λ_em_= 613 nm). The excitation bandwidth was 9 nm. The emission bandwidth was 20 nm. The gain was set to 115. The image generated with the repressor encoding genetic cascade was additionally collected with the Spinach aptamer (λ_ex_= 469 nm, λ_em_= 501 nm).

**Protein and RNA Standard Curves**: Standard curves to translate fluorescence intensity into molar concentrations were generated by using recombinantly expressed and purified fluorescent proteins. His-tagged versions of GFPmut3b, Azurite, and mRFP1 were generated by mutating the stop codon within pET21b. The resulting constructs contained an additional 24 residues including a carboxy-terminal hexahistidine-tag. Each His-tagged construct was purified as described previously.^16^ Similarly, purified RNA harboring the Spinach aptamer and transcribed from the different constructs was used to equate the fluorescence intensity of the Spinach aptamer with molar concentrations. Transcription reactions were assembled and then purified as previously described.^16^

**Computational kinetic model**: COPASI^50^ was used to implement a computational model that consisted of 10 reactions and 12 parameters describing cell-free RNA and protein expression (Figure S10, Table S6, Table S7, Table S8). The resources necessary for transcription and translation were modeled as a single species (R1 and R2, respectively) and were subject to degradation. Here, resources refer not only to the molecular machinery required to sustain transcription and translation, such as the T7 RNA polymerase, the ribosomes and all the accessory protein factors, but also to the small molecules required to sustain each reaction. Following the PURE system composition,^17^ only the initial concentration of T7 RNA polymerase and ribosomes were defined. For the rest of the species and for the reaction rates, parameter estimations based on experimental data were performed. Parameter estimations were also included for two noise parameters, including a ±5% range for the template DNA concentration and a ±10% range for the composition of the PURE system.

**Statistical analysis**: The coefficient of variation was calculated as the ratio between the standard deviation and the mean value of each experiment (C_v_=σ/μ). The coefficient of variation for single batch experiments was calculated by averaging the coefficients of variation of three replicates, each collected with the same batch of the PURE system. The coefficient of variation for multiple batch experiments was calculated by averaging the coefficients of variation of nine replicates, using three different PURE system batches (three replicates per batch).

**Photobleaching**: 20 μL of 2.5 μM RFP, 2.5 μM BFP, and 2.5 μM GFP purified protein solutions in 10 mM Tris-HCl, pH 7.5 were loaded into a Greiner Flat Bottom Black Polystyrene 384-well plate (pre-heated for 10 min at 37 °C). A Tecan Infinite PRO200 plate reader was used to record the fluorescence of the single proteins with their specific excitation settings followed by measurements at each of the following wavelengths in the order listed: Blue (λ^ex^= 377 nm, λ^em^= 472 nm), Red (λ^ex^= 579 nm, λ^em^= 613 nm), Green (λ^ex^= 474 nm, λ^em^= 511 nm). The excitation bandwidth was 9 nm and the emission bandwidth was 20 nm. The gain was set to 115.

### Supporting Information

Supporting Information Available: Tables S1-S17 and Figures S1-S21. Links to the deposited genetic sequences in the ACS Synthetic Biology Registry. This material is available free of charge via the Internet at http://pubs.acs.org.

## AUTHOR INFORMATION

CIBIO, University of Trento, via Sommarive 9, 38123 Povo (TN), Italy

## ACKNOWLEDGMENTS

We thank the Armenise-Harvard foundation and CIBIO for funding, Cristina Del Bianco of the protein technology facility at CIBIO for assistance, and Federico Brunello for help with cloning and the initial characterization of BFP. This research was funded in part by the Autonomous Province of Trento, “Grandi Progetti 2012”, project “Characterizing and improving brain mechanisms of attention-ATTEND”.

